# Comparison of different models in predicting habitat suitability of rare species in Uzbekistan: 8 rare Tulipa species case-study

**DOI:** 10.1101/2022.07.08.499262

**Authors:** Khondamir Rustamov

## Abstract

Species distribution models (SDMs) have become an essential tool in conservational biology, biogeography and ecology. But there is no consequence in what SDM method is the most efficient in predicting suitable habitat distribution of rare species. To explore this issue, we chose 8 rare *Tulipa* species in Uzbekistan as case study to test 8 common Machine Learning (GLM, GBM, MARS, CTA, SRE, FDA, RF, MaxEnt) and Deep Neural Network (DNN) SDM models, using three different methods of pseudo-absence data generation (random sampling, random sampling with exclusion buffer, random sampling with environmental profiling). To compare the effectiveness of each model 3 common metrics (Area under ROC (AUC), True skill statistics (TSS) and Cohen’s Kappa (K)) were used. We have found that RF and GBM combined with RSEP strategy are superior to other modeling methods.

## Introduction

Species Distribution Models (SDMs) are powerful and efficient tool for predicting potential habitat distribution for species, so have a significant impact in conservational biology and ecology (Guisan and Zimmermann, 2000; Elith and Leathwick, 2009; McCune, 2016). Using SDMs for studying rare plant species in Uzbekistan is in the initial stage now. Only a few studies were carried out (Baikov et al. 2021; Khujanov, 2021; Mavlanov et al. 2021) and most of them were conducted using MaxEnt (Philips et al. 2006), which is considered the most powerful machine learning algorithm with an uncomplicated graphical user interface, which made MaxEnt the most comfortable and popular technique for researchers in last years (Philips et al. 2017). However, numerous studies demonstrated that Random Forest and other machine learning techniques in some cases are more efficient than MaxEnt (Williams et al. 2009; Duan et al. 2014; Mi et al. 2017).

Furthermore, recent works illustrated that different deep learning models, such as convolutional neural networks (Deneu et al. 2021) and deep neural networks (Rew et al. 2021; Seo et al. 2021), gain higher results than traditional machine learning algorithms in predicting species distribution.

Most machine learning models require presence and absence data as well. It’s problematic to collect real absence data for rare species, so pseudo-absence data is generated. There are few approaches to generating pseudo-absence data (Chefaoui and Lobo, 2008; Iturbide et al. 2015). In this study, three methods of pseudo absence generating (Random Sampling (RS), Random Sampling with Exclusion Buffer (RSEB), and Random Sampling with Ecological Profiling (RSEP)) were tested.

In this research we estimated the efficiency of different modeling algorithms in bioclimatic conditions of Uzbekistan using AUC, TSS and Cohen’s Kappa metrics. To perform that we chose 8 rare Tulipa species (*T. kaufmanniana, T. ferganica, T. dubia, T. affinis, T. tubergeniana, T. fosteriana, T. carinata, T. mogoltavica*) distributed in Central Asian Mountain botanical-geographical province of Uzbekistan to evaluate nine SDMs (DNN, GLM, GBM, RF, MARS, CTA, SRE, FDA, MaxEnt) and three pseudo-absence generating strategies (RS, RSEB, RSEP).

## Materials and Methods

### Dataset construction

8 rare *Tulipa* species with different distribution patterns and number of records were chosen to conduct research (Table 1). The true presence data of target species was collected from appropriate literature (Tojibaev and Beshko, 2015; Dekhkonov et al. 2021).

**Table 1.** Target species characteristics.

| № | Species name | Number of records | IUCN category | Uzbek RDB category (2019) |
| --- | --- | --- | --- | --- |
| 1 | *T. kaufmanniana* Regel | 45 | NT | 3 (Reducing) |
| 2 | *T. ferganica* Vved. | 37 | VU B2b (ii, iii, iv) | 2 (Rare) |
| 3 | *T. dubia* Vved. | 33 | VU, B1b (ii, iii, iv) 2b (ii, iii, iv) | 3 (reducing) |
| 4 | *T. affinis* Botschanz. | 19 | VU, B1b (ii, iii, iv) 2b (ii, iii, iv) | 3 (reducing) |
| 5 | *T. tubergeniana* Hoog. | 15 | EN A2cd B1b(ii, iii, iv) 2b(ii, iii, iv) | 3 (reducing) |
| 6 | *T. fosteriana* Hoog ex W. Irving | 12 | EN B1b(ii, iii, iv) 2b(ii, iii, iv) | 2 (Rare) |
| 7 | *T. carinata* Vved. | 17 | VU B1b(ii, iii, iv) 2b(ii, iii, iv) | 3 (reducing) |
| 8 | *T. mogoltavica* Popov & Vved. | 15 | VU B1b(ii, iii, iv) 2b(ii, iii, iv) | - |

We used 19 WordClim environmental variables and attitude data (Table 2) at 30-s spatial resolution in a GIS format to extract predictor values for Central Asian Mountain province of Uzbekistan, it lies approximately between latitudes 36º to 44º N and longitudes 65º to 75º E (Fick and Hijmans, 2017).

**Table 2.** List of 20 environmental variables.

| Variable | Description |
| --- | --- |
| **BIO1** | Annual Mean Temperature |
| **BIO2** | Mean Diurnal Range (Mean of monthly (max temp - min temp)) |
| **BIO3** | Isothermality (BIO2/BIO7) (×100) |
| **BIO4** | Temperature Seasonality (standard deviation ×100) |
| **BIO5** | Max Temperature of Warmest Month |
| **BIO6** | Min Temperature of Coldest Month |
| **BIO7** | Temperature Annual Range (BIO5-BIO6) |
| **BIO8** | Mean Temperature of Wettest Quarter |
| **BIO9** | Mean Temperature of Driest Quarter |
| **BIO10** | Mean Temperature of Warmest Quarter |
| **BIO11** | Mean Temperature of Coldest Quarter |
| **BIO12** | Annual Precipitation |
| **BIO13** | Precipitation of Wettest Month |
| **BIO14** | Precipitation of Driest Month |
| **BIO15** | Precipitation Seasonality (Coefficient of Variation) |
| **BIO16** | Precipitation of Wettest Quarter |
| **BIO17** | Precipitation of Driest Quarter |
| **BIO18** | Precipitation of Warmest Quarter |
| **BIO19** | Precipitation of Coldest Quarter |
| **Attitude** | Attitude |

### Pseudo-Absence Generation

To generate effective pseudo-absence data for target species we investigated different approaches. The simplest method of pseudo-absence data generation is Random Sampling (RS), according to this method pseudo-absences are randomly sampled from all study area without any limitations. Whereas Random Sampling with Exclusion Buffer (RSEB) randomly chooses points that are not within some minimum selected distance to any presence point, this distance is referred to as Exclusion Buffer. Several researchers recommended setting a 10 km exclusion buffer to avoid collecting presence and pseudo-absence at the same grids. We also use a 10 km exclusion buffer to generate pseudo-absence data by the RSEB technique. Random Sampling with Environmental Profiling (RSEP) tries to define the possible area in which presence records can occur and select pseudo-absences out of this area. RSEP can be calculated using One-Class Support Vector Machine (OCSVM). We generated 10 times more pseudo-absence data than presence data by three sampling methods using Python programming language, moreover, Scikit-learn Python library was used to provide RSEP (Pedregosa et al. 2011). The attached pseudo-absence generating code is given in the Supplementary Materials. Fig. 1 compares pseudo-absence data generated by RS, RSEB, RSEP for *T. kaufmanniana*.

**Fig. 1.**
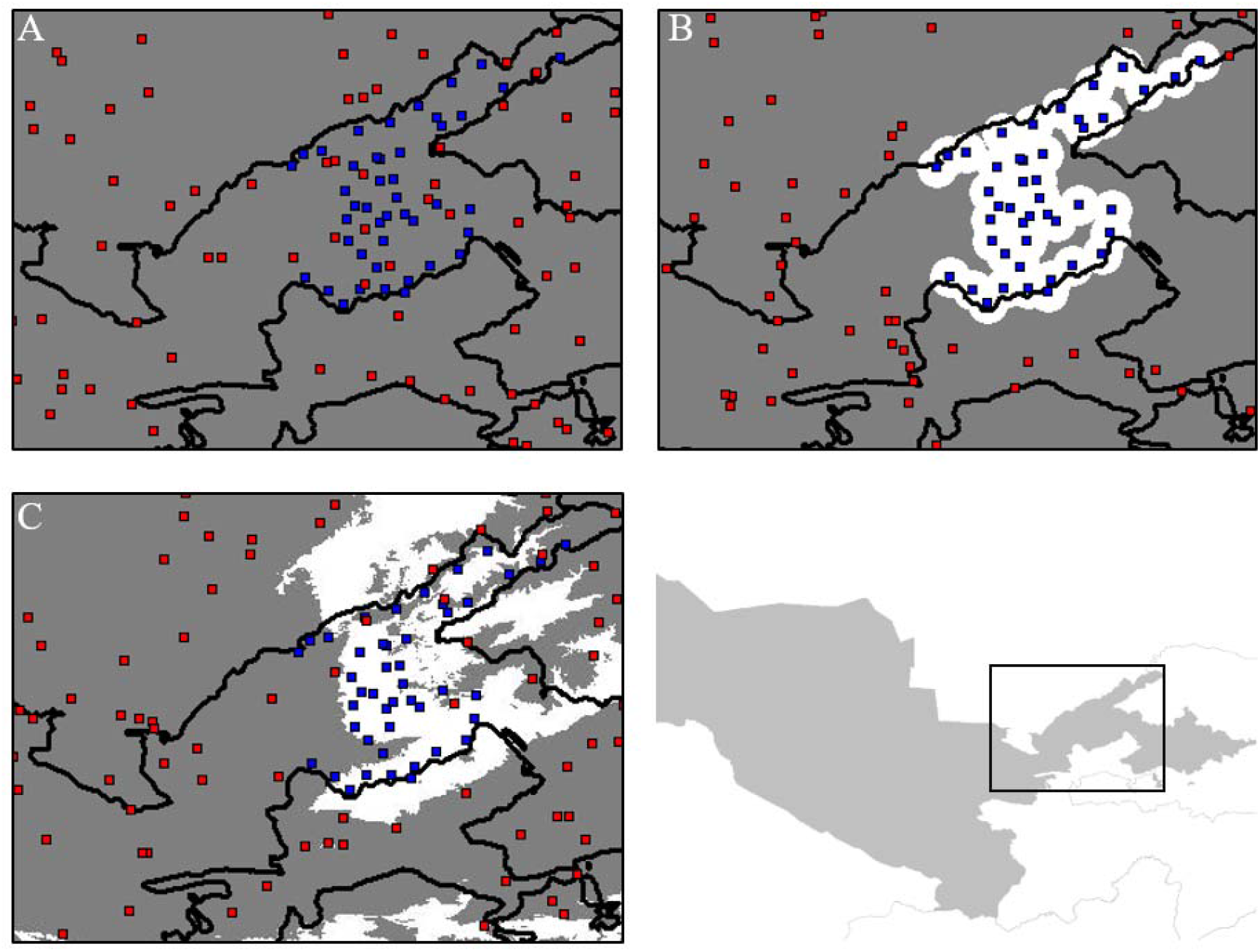
Different methods of generating pseudo-absence data for *T. kaufmanniana*. A – random sampling (RS); B – random sampling with exclusion buffer (RSEB), white circles illustrate 10 km exclusion buffer; C – random sampling with ecological profiling (RSEP) with OCSVM prediction map.

### Species distribution models

We fitted SDMs using 8 machine learning approaches implemented in the ‘BIOMOD2’ R package (Thuiller et al. 2009). These were generalized linear models (GLM), generalized boosting models (GBM), random forest (RF), multivariate adaptive regression splines (MARS), classification tree analysis (CTA), surface range envelop (SRE), flexible discriminant analysis (FDA), maximum entropy (MaxEnt). We validated the predictions of our models by using five-fold cross-validation with five-time evaluation runs and recording their average results. Records were split at an 80%-20% ratio of training and testing data. Table 3 illustrates other training strategies and selected variables for all 8 models.

**Table 3:** Training strategies and selected parameters for SDMs

| Model | Training strategies and parameters | Software |
| --- | --- | --- |
| **Deep Neural Network (DNN)** | 3 hidden layers (with 200, 300, 300 neurons each), dropout, 1000 epochs with early stopping, rectified linear unit (ReLU) activation, Adam optimization with learning rate 0.001 | Tensorflow (Python) |
| **Generalized Linear Model (GLM)** | Quadratic regression. Stepwise procedure using Akaike Information criteria | Biomod2 (R) |
| **Generalized Boosting Model (GBM)** | Bernoulli distribution. 2500 trees, depth 7, 5 terminal nodes, learning rate 0.0001 | Biomod2 (R) |
| **Random Forest (RF)** | 500 trees, 5 nodes (Biomod2 auto-optimization) | Biomod2 (R) |
| **Multiple Adaptive Regression Splines (MARS)** | Simple piecewise linear, threshold = 0.001, backward pruning | Biomod2 (R) |
| **Classification Tree Analysis (CTA)** | Categorical classification (Biomod2 auto-optimization) | Biomod2 (R) |
| **Surface Range Envelop (SRE)** | 0.025 quantile for extreme environmental variable | Biomod2 (R) |
| **Flexible Discriminant Analysis (FDA)** | MARS method | Biomod2 (R) |
| **Maximum Entropy (MaxEnt)** | 200 iterations, linear and quadratic variables (Biomod2 auto-optimization) | Biomod2 (R) |

### DNN model construction

We used the TensorFlow Python library to build and train the DNN model. Our DNN model receives 20 WordClim variables on the input layer and perform it at hidden layers to produce the final prediction on the output layer, which is represented two values, regarding as probabilities of species absence and presence at given bioclimatic inputs. The structure of the DNN model is illustrated on Fig. 2.

**Fig. 2.**
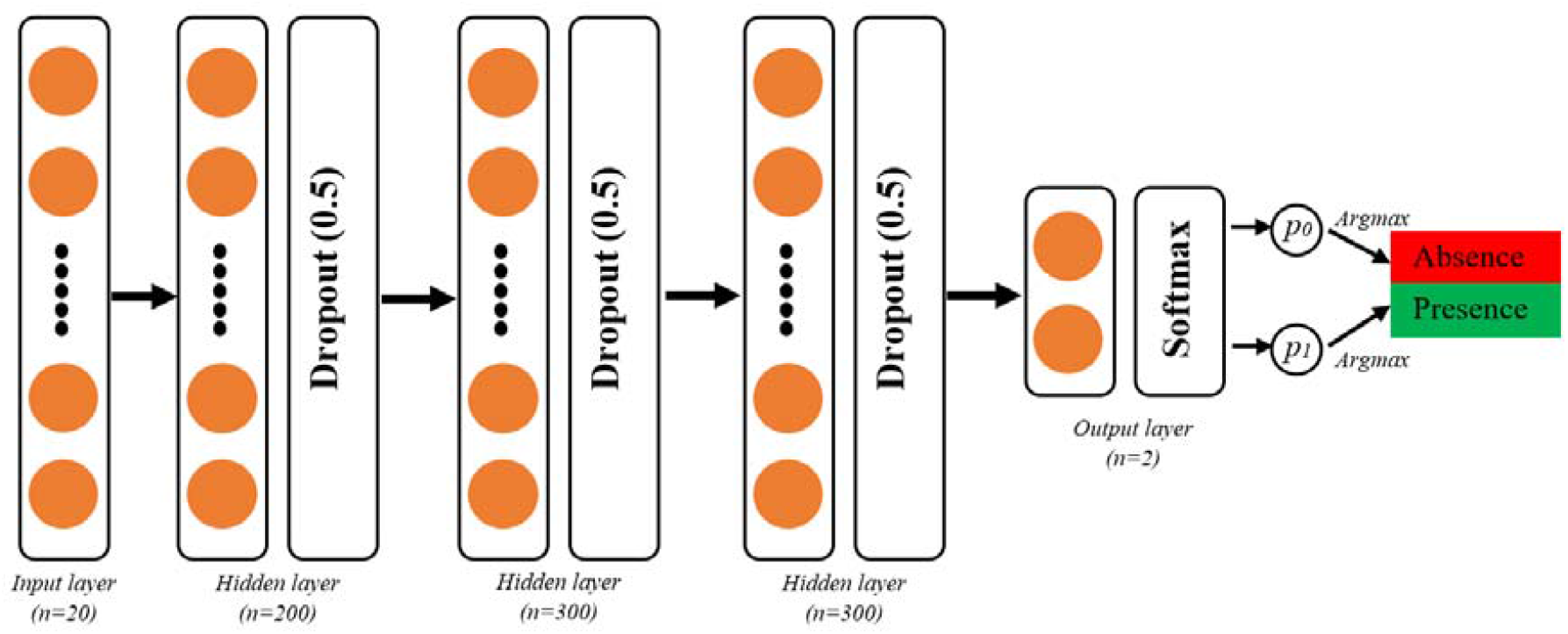
The structure of Deep Neural Network (DNN) model.

To optimize the model, we used three hidden layers (with 200, 300, and 300 neurons each), a batch size equal to 1/10 of training data, and 1000 epochs with an early stopping callback. Adaptive moment estimation (Adam) with a learning rate of 0.001 was used to update model weights iterative based on training data, rectified linear unit (ReLU) activation function to deal with the vanishing gradient problem, and categorical cross entropy loss function to quantify the difference between two classes.

We split data into three groups for our DNN performing (70% - training, 10% - validation, and 20% test data). As pseudo-absence data was 10 times bigger than presence data our dataset was imbalanced. To deal with the imbalanced data classification problem we set different class weights for presence and pseudo-absence classes,

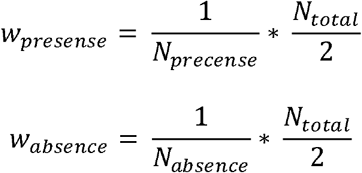

where, *w*_*presence*_ and *w*_*absence*_ are weights for presence and pseudo-absence classes, *N*_*total*_ – number of total samplings, *N*_*presence*_ and *N*_*absence*_ are number of presence and pseudo-absence records respectively. Scaling by *N*_*total*_*/2* helps keep the loss to a similar magnitude.

### Evaluation approach

In this research three common evaluation metrics were used: area under curve (AUC), true skill statistics (TSS) and Kappa statistics (K). Scikit-learn Python package (Pedregosa et al. 2011) was used to determine AUC and confusion matrix to calculate both K and TSS. To define true positives (TP), false positives (FP), true negatives (TN) and false negatives (FN) in confusion matrix we selected threshold that maximized both the True Positive Rate (TPR) and True Negative Rate (TNR) – Max (Specificity + Sensitivity) threshold optimization.

Model, which performs AUC and K ≥ 0.9, TSS ≥ 0.8 is excellent; AUC and K between 0.8 – 0.9 and TSS between 0.6 – 0.8 = good; AUC, K between 0.7 – 0.8 and TSS between 0.4 – 0.6 = fair; while AUC ≤ 0.7, K ≤ 0.6 and TSS ≤ 0.4 is considering as poor and without predictive ability.

### Prediction maps

The most defined models produce not binary (0 or 1), but continuous prediction maps. To estimate the possible range of species distribution we have to set a threshold value below which a species is considered absent. To perform these calculations, we used Max (Specificity + Sensitivity) threshold optimizing strategy, which is also known as TSS threshold optimizing, to generate binary prediction maps for all target species.

## Results

### SDMs performance

We tested the efficiency for 9 different SDM approaches using 3 various pseudo-absence generating strategies. Table 4 illustrates the average performance of different SDM techniques for three pseudo-absence sampling techniques. Prediction maps for all species using all models and RSEP pseudo-absence selecting strategy are illustrated in Figs. 3-10. Table 5 illustrates the suitable distribution area for all target species calculated by using all models and RSEP pseudo-absence generating technique. All SDMs results for 8 Tulipa species using 3 pseudo-absence generating strategies are showed in supplementary materials.

**Table 4.** Average metrics (AUC, TSS, Kappa) for all models among 8 species.

| | AUC (Mean ± SD) | | | TSS (Mean ± SD) | | | Kappa (Mean ± SD) | | |
| --- | --- | --- | --- | --- | --- | --- | --- | --- | --- |
|  | RS | RSEB | RSEP | RS | RSEB | RSEP | RS | RSEB | RSEP |
| **DNN** | 0.969±<br>0.0169 | 0.977±<br>0.0238 | 0.991±<br>0.0097 | 0.918±<br>0.0493 | 0.943±<br>0.0432 | 0.961±<br>0.0444 | 0.767±<br>0.1023 | 0.829±<br>0.133 | 0.891±<br>0.1349 |
| **GLM** | 0.979±<br>0.0179 | 0.984±<br>0.0283 | 0.982±<br>0.027 | 0.916±<br>0.0673 | 0.948±<br>0.0805 | 0.913±<br>0.0818 | 0.802±<br>0.149 | 0.876±<br>0.1731 | 0.844±<br>0.1621 |
| **GBM** | 0.999±<br>0.0012 | 0.999±<br>0.0003 | 0.999±<br>0.0003 | 0.988±<br>0.0096 | 0.995±<br>0.0039 | 0.996±<br>0.0047 | 0.97±<br>0.0276 | 0.982±<br>0.0184 | 0.985±<br>0.0164 |
| **RF** | 1.000±<br>0.0 | 0.999±<br>0.0003 | 0.999±<br>0.0004 | 0.996±<br>0.0024 | 0.997±<br>0.0027 | 0.998±<br>0.0032 | 0.982±<br>0.0128 | 0.988±<br>0.0134 | 0.99±<br>0.0174 |
| **MARS** | 0.985±<br>0.0248 | 0.987±<br>0.0225 | 0.998±<br>0.0014 | 0.954±<br>0.0525 | 0.960±<br>0.0518 | 0.988±<br>0.0167 | 0.902±<br>0.0837 | 0.934±<br>0.0725 | 0.977±<br>0.0313 |
| **CTA** | 0.945±<br>0.024 | 0.914±<br>0.0949 | 0.918±<br>0.0889 | 0.825±<br>0.0779 | 0.820±<br>0.1717 | 0.771±<br>0.1562 | 0.8±<br>0.0392 | 0.82±<br>0.1431 | 0.798±<br>0.1351 |
| **SRE** | 0.789±<br>0.0559 | 0.794±<br>0.0574 | 0.796±<br>0.0598 | 0.578±<br>0.1119 | 0.588±<br>0.1149 | 0.593±<br>0.1198 | 0.629±<br>0.07 | 0.672±<br>0.0818 | 0.693±<br>0.1029 |
| **FDA** | 0.983±<br>0.0117 | 0.983±<br>0.0175 | 0.982±<br>0.0135 | 0.935±<br>0.0454 | 0.944±<br>0.0419 | 0.951±<br>0.0302 | 0.85±<br>0.0895 | 0.909±<br>0.0689 | 0.927±<br>0.0439 |
| **MaxEnt** | 0.922±<br>0.0562 | 0.937±<br>0.0439 | 0.946±<br>0.0436 | 0.834±<br>0.1054 | 0.876±<br>0.0871 | 0.893±<br>0.0867 | 0.854±<br>0.0625 | 0.908±<br>0.0504 | 0.886±<br>0.0868 |

*SD is an abbreviation for Standard Deviation*

**Fig. 3.**
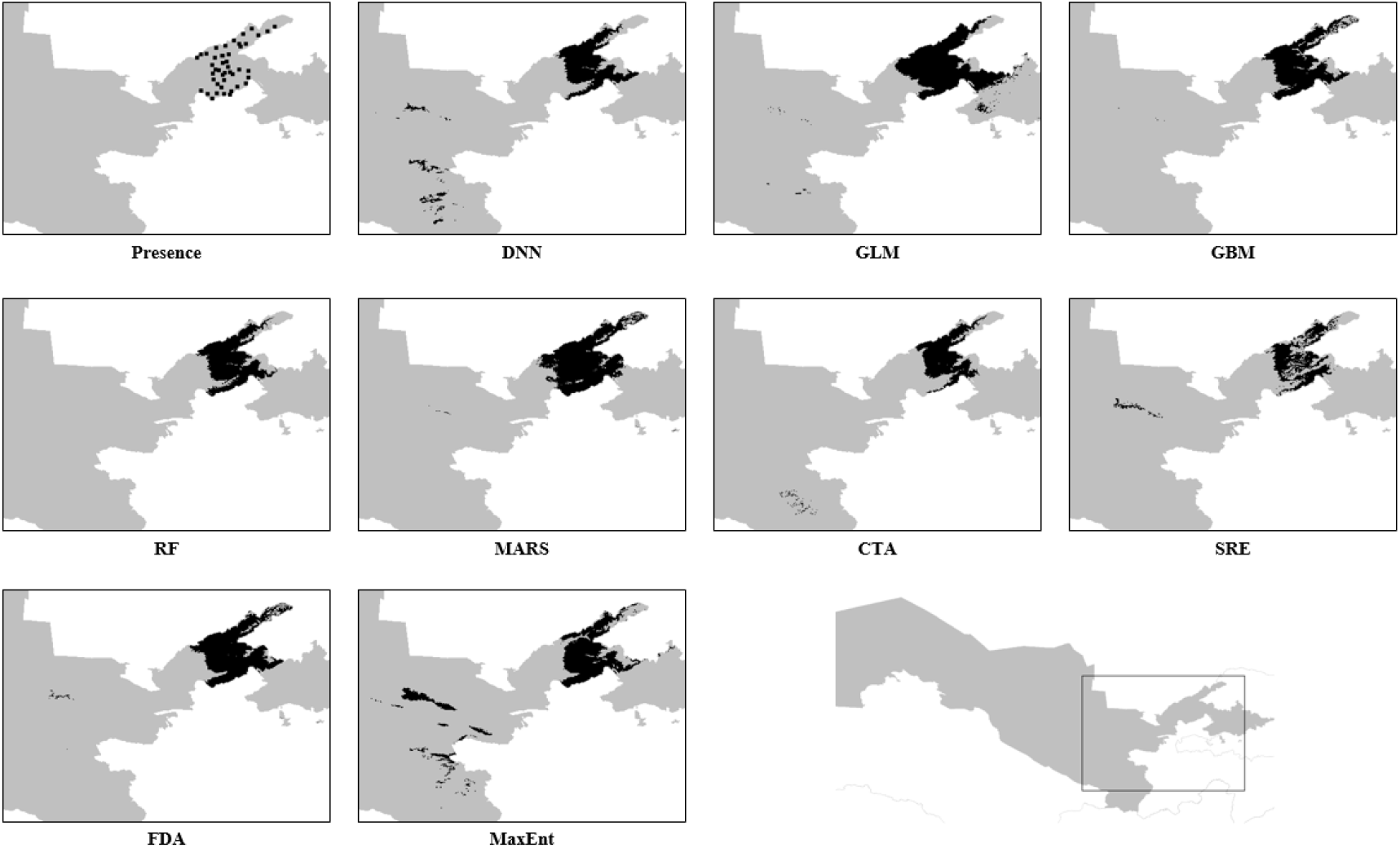
Prediction maps for *T. kaufmanniana* in nine SDMs (DNN, GLM, GBM, RF, MARS, CTA, SRE, FDA and MaxEnt) by using RSEP pseudo-absence generating strategy.

**Fig. 4.**
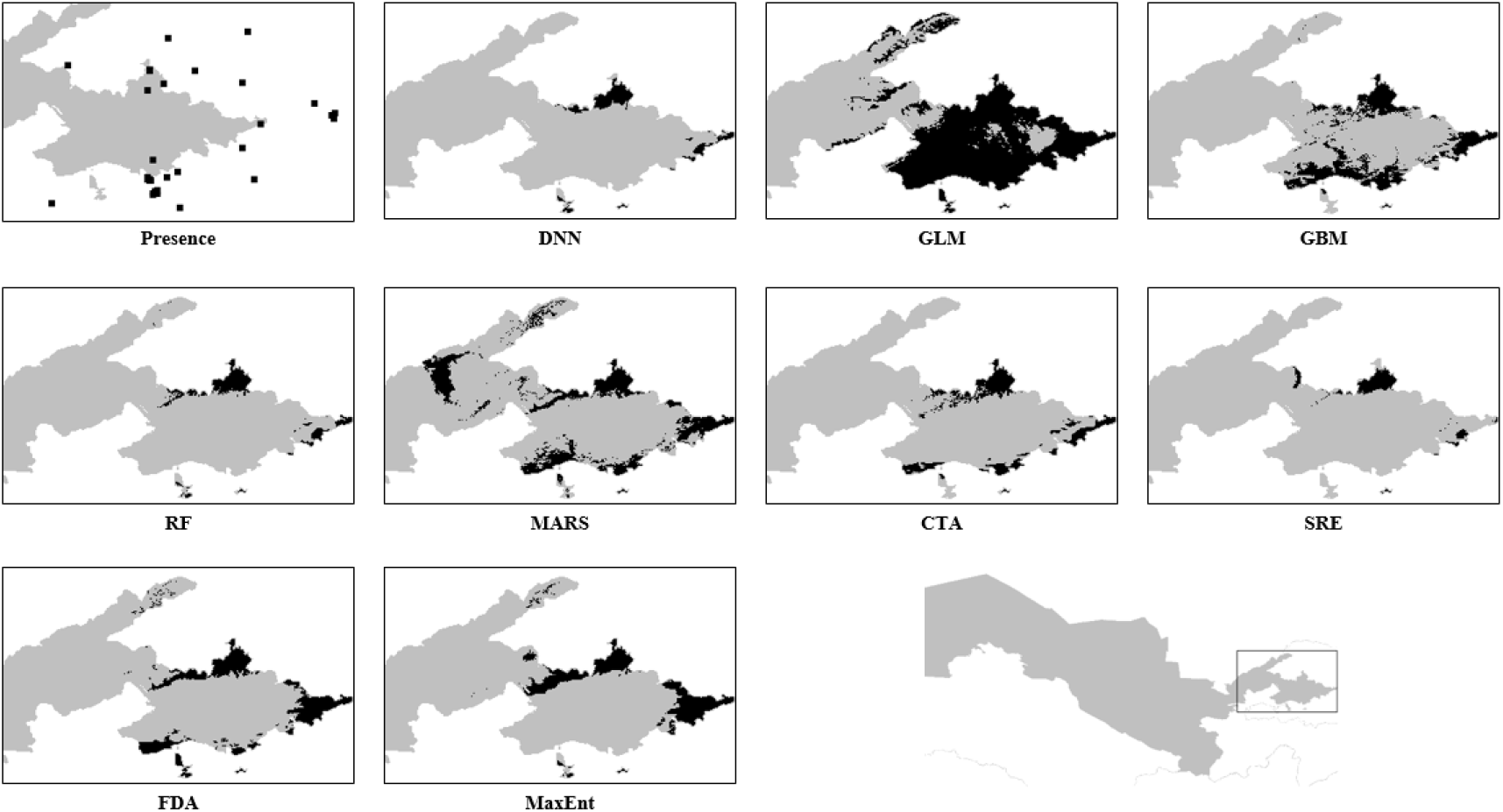
Prediction maps for *T. ferganica* in nine SDMs (DNN, GLM, GBM, RF, MARS, CTA, SRE, FDA and MaxEnt) by using RSEP pseudo-absence generating strategy.

**Fig. 5.**
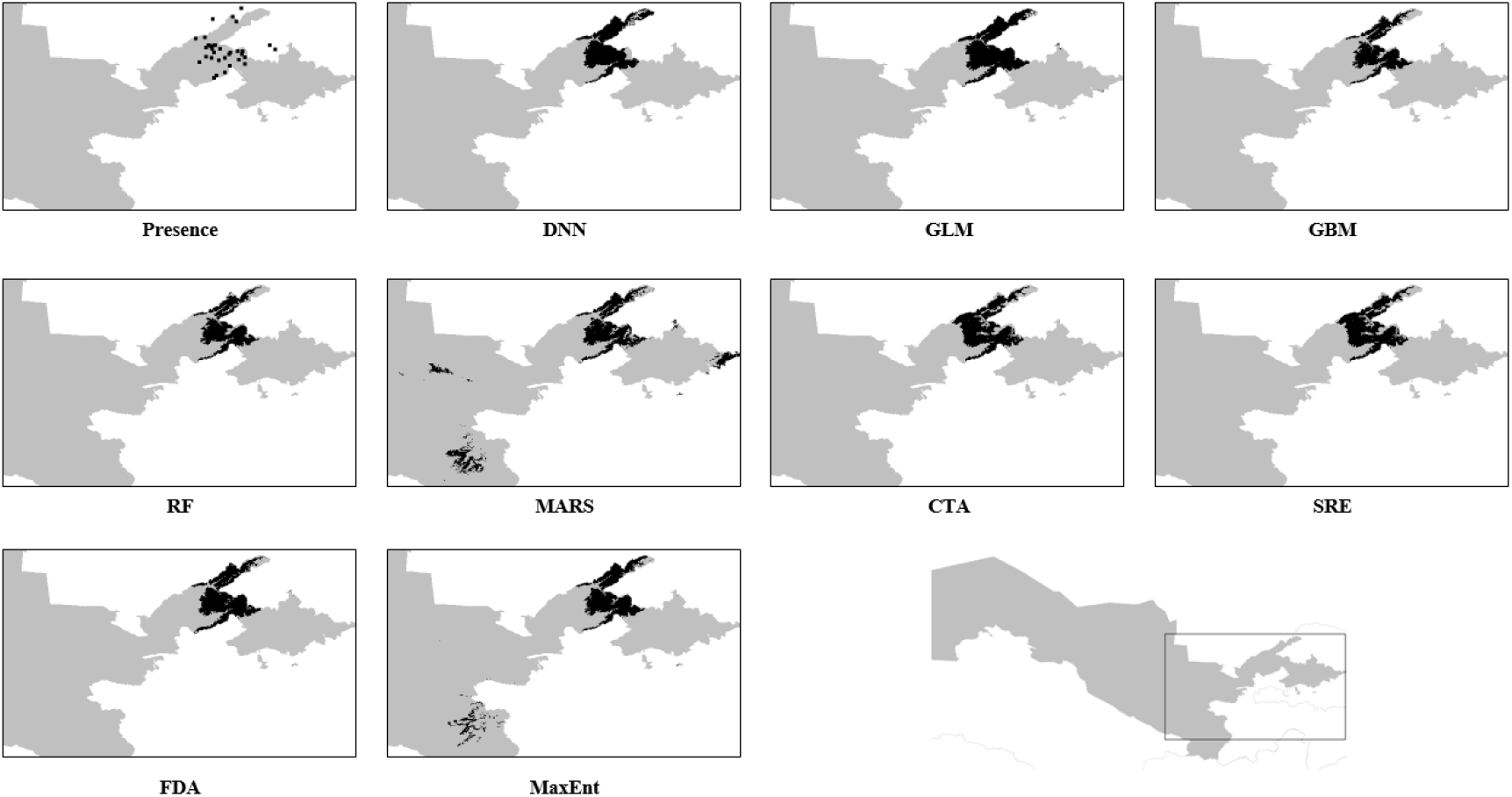
Prediction maps for *T. dubia* in nine SDMs (DNN, GLM, GBM, RF, MARS, CTA, SRE, FDA and MaxEnt) by using RSEP pseudo-absence generating strategy.

**Fig. 6.**
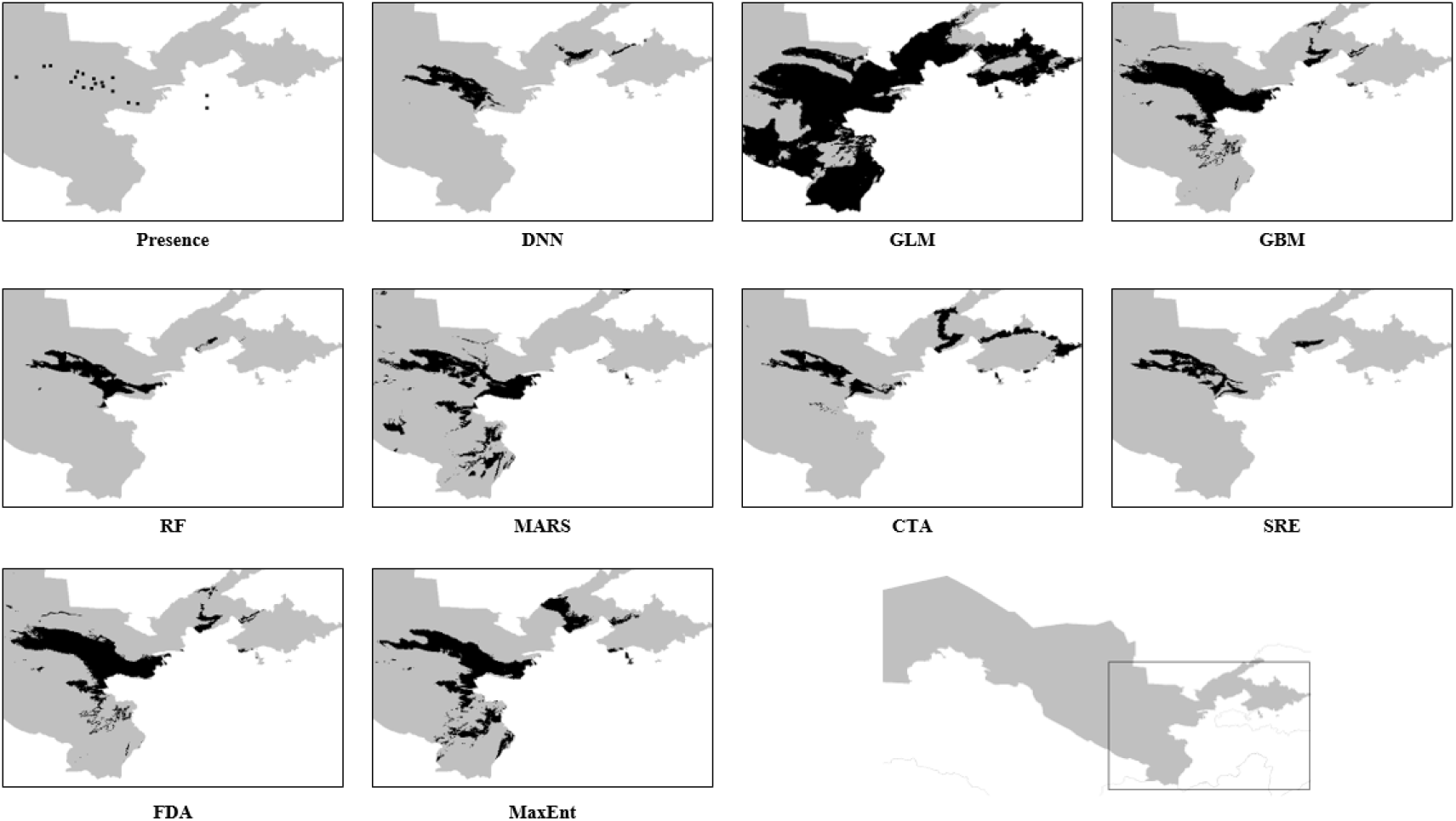
Prediction maps for *T. affinis* in nine SDMs (DNN, GLM, GBM, RF, MARS, CTA, SRE, FDA and MaxEnt) by using RSEP pseudo-absence generating strategy.

**Fig. 7.**
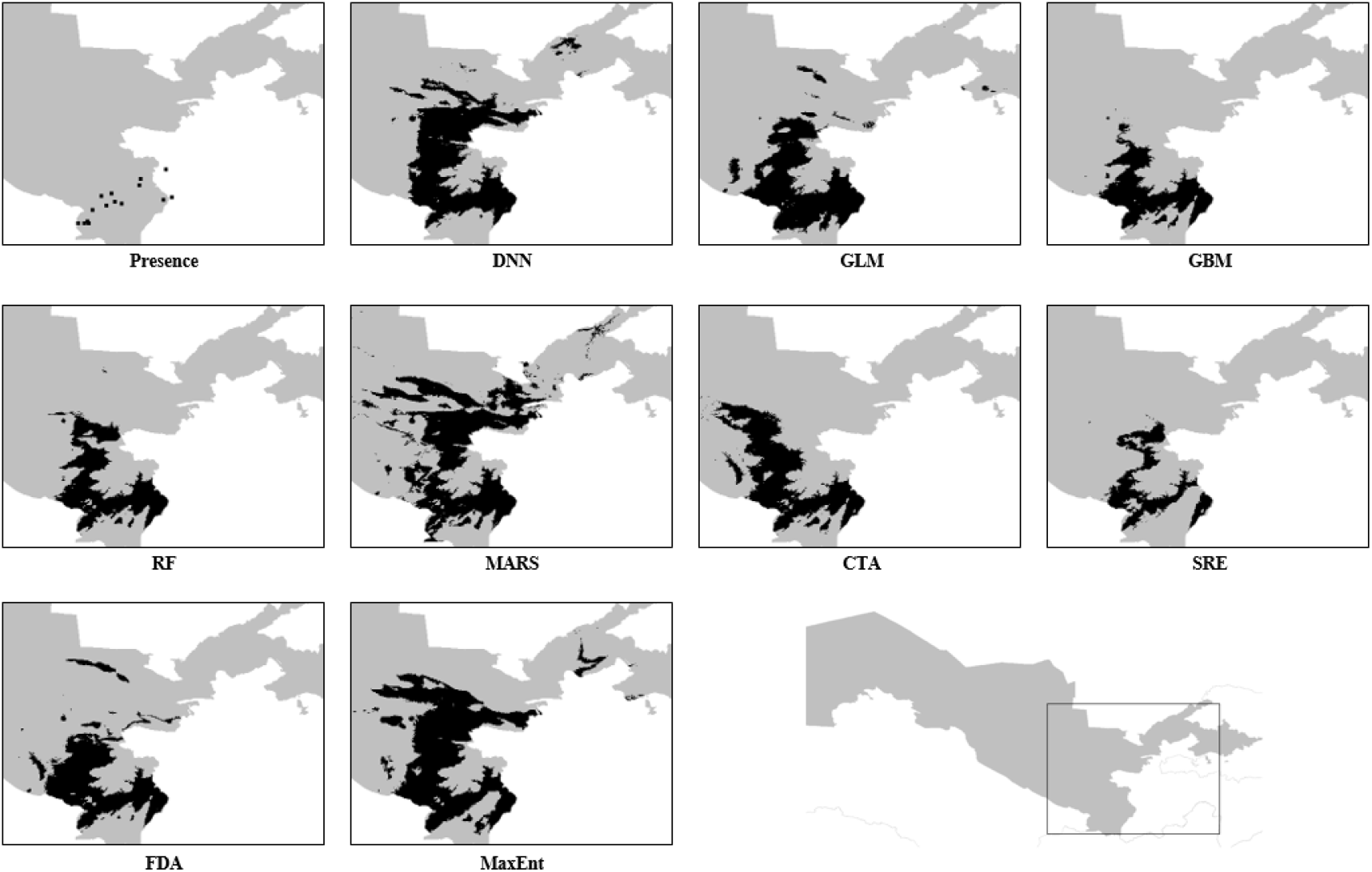
Prediction maps for *T. tubergeniana* in nine SDMs (DNN, GLM, GBM, RF, MARS, CTA, SRE, FDA and MaxEnt) by using RSEP pseudo-absence generating strategy.

**Fig. 8.**
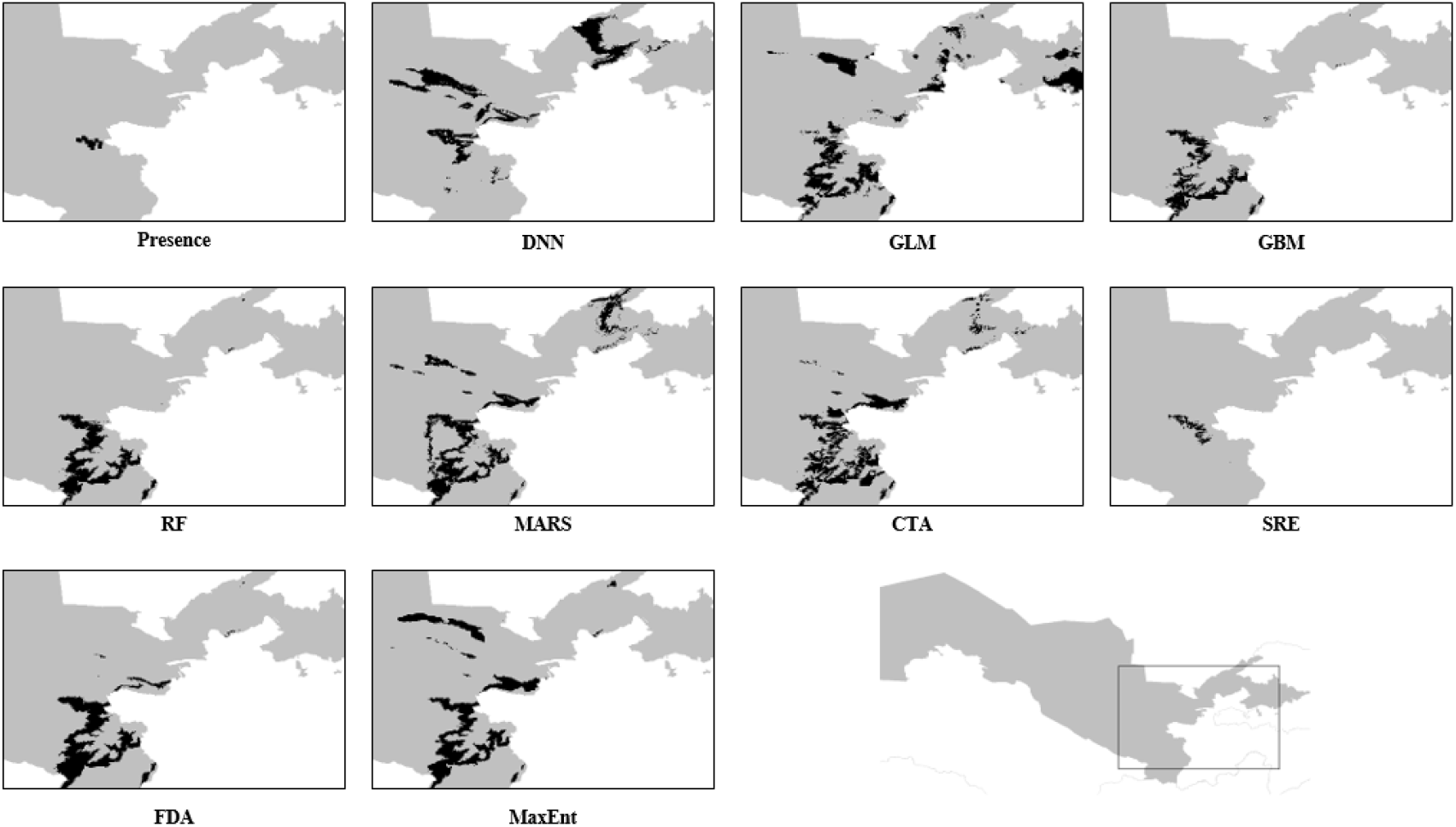
Prediction maps for *T. fosteriana* in nine SDMs (DNN, GLM, GBM, RF, MARS, CTA, SRE, FDA and MaxEnt) by using RSEP pseudo-absence generating strategy.

**Fig. 9.**
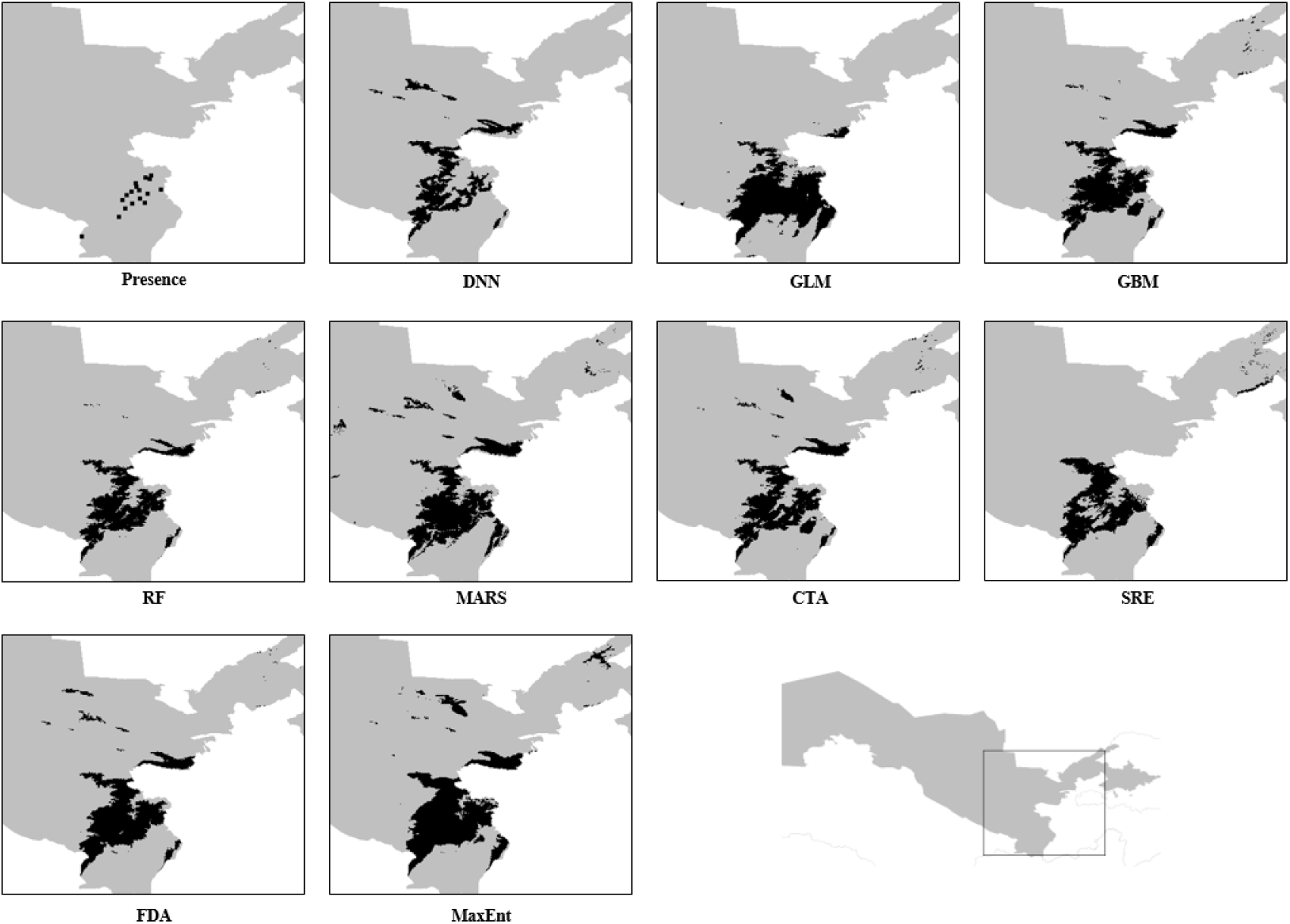
Prediction maps for *T. carinata* in nine SDMs (DNN, GLM, GBM, RF, MARS, CTA, SRE, FDA and MaxEnt) by using RSEP pseudo-absence generating strategy.

**Fig. 10.**
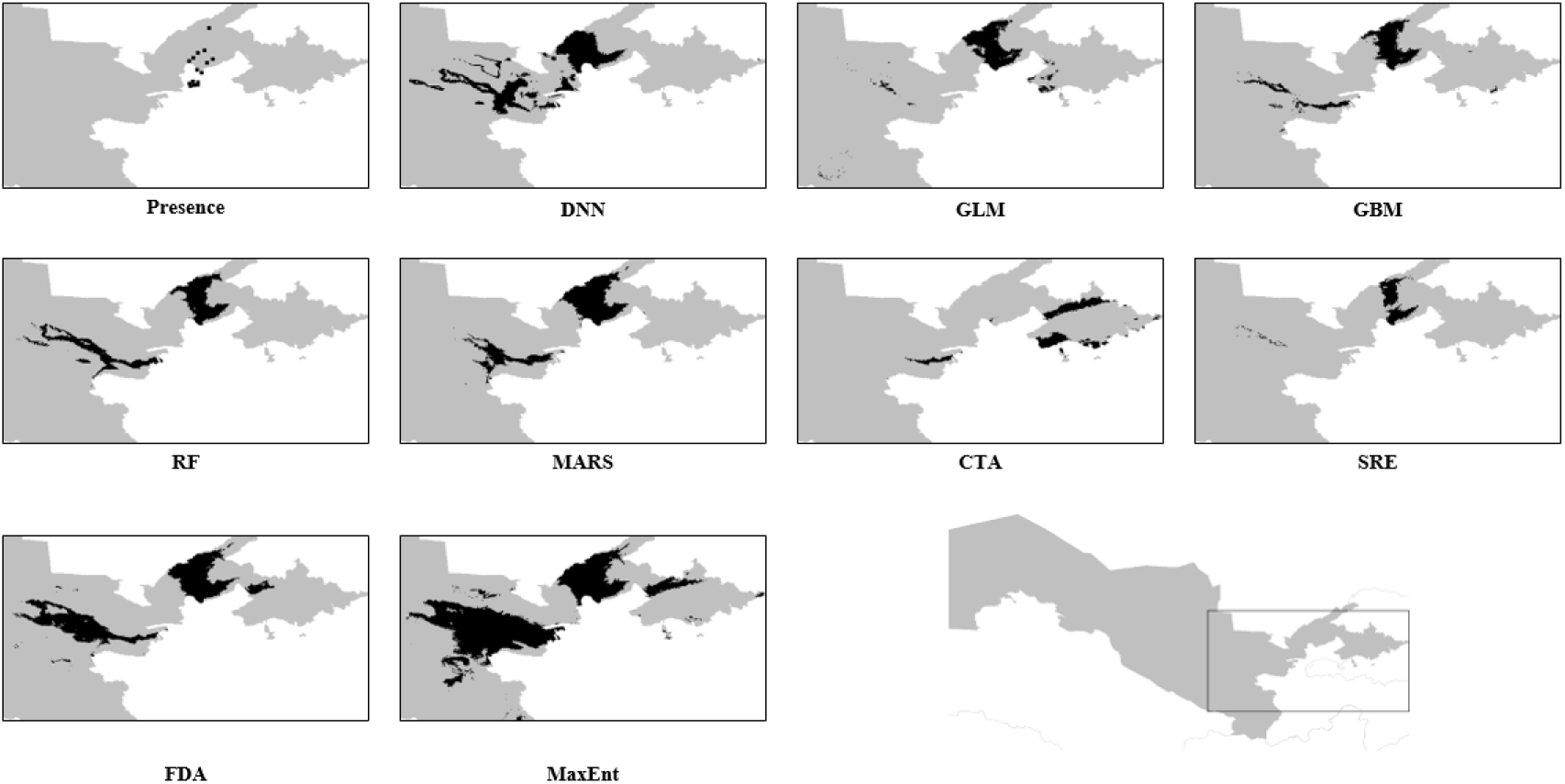
Prediction maps for *T. mogoltavica* in nine SDMs (DNN, GLM, GBM, RF, MARS, CTA, SRE, FDA and MaxEnt) by using RSEP pseudo-absence generating strategy.

**Table 5.** Suitable distribution area (km^2^) for all 8 Tulipa species in Uzbekistan using all models and RSEP pseudo-absence generating strategy.

|  | *T. kauf* | *T. ferg* | *T. dubia* | *T. affinis* | *T. tuberg* | *T. fost* | *T. car* | *T. mog* |
| --- | --- | --- | --- | --- | --- | --- | --- | --- |
| **DNN** | 8365<br>(1.87%) | 1276<br>(0.285%) | 6677<br>(1.492%) | 7420<br>(1.658%) | 30646<br>(6.85%) | 9447<br>(2.111%) | 9098<br>(2.034%) | 11787<br>(2.635%) |
| **GLM** | 12137<br>(2.713%) | 15981<br>(3.572%) | 8018<br>(1.792%) | 90988<br>(20.337%) | 24651<br>(5.51%) | 11890<br>(2.658%) | 14722<br>(3.291%) | 6002<br>(1.342%) |
| **GBM** | 8765<br>(1.959%) | 4946<br>(1.106%) | 5761<br>(1.288%) | 23033<br>(5.148%) | 14855<br>(3.32%) | 5289<br>(1.182%) | 12240<br>(2.736%) | 6101<br>(1.364%) |
| **RF** | 7728<br>(1.727%) | 1514<br>(0.338%) | 5642<br>(1.261%) | 8910<br>(1.991%) | 17517<br>(3.915%) | 6757<br>(1.51%) | 10446<br>(2.335%) | 8428<br>(1.884%) |
| **MARS** | 11093<br>(2.479%) | 6047<br>(1.352%) | 8934<br>(1.997%) | 18274<br>(4.084%) | 35333<br>(7.897%) | 10736<br>(2.4%) | 14635<br>(3.271%) | 9538<br>(2.132%) |
| **CTA** | 6225<br>(1.391%) | 2671<br>(0.597%) | 7100<br>(1.587%) | 12730<br>(2.845%) | 23245<br>(5.196%) | 9595<br>(2.145%) | 11804<br>(2.638%) | 4085<br>(0.913%) |
| **SRE** | 5841<br>(1.305%) | 1074<br>(0.24%) | 4112<br>(0.919%) | 7294<br>(1.63%) | 10845<br>(2.424%) | 814<br>(0.182%) | 10725<br>(2.397%) | 2700<br>(0.603%) |
| **FDA** | 10099<br>(2.257%) | 3962<br>(0.886%) | 6952<br>(1.554%) | 23033<br>(5.148%) | 21605<br>(4.829%) | 9019<br>(2.016%) | 13531<br>(3.024%) | 14554<br>(3.253%) |
| **MaxEnt** | 10280<br>(2.298%) | 3760<br>(0.84%) | 7522<br>(1.681%) | 23269<br>(5.201%) | 36985<br>(8.267%) | 10691<br>(2.39%) | 17128<br>(3.828%) | 27840<br>(6.223%) |

*Abbreviations: T. kauf – T. kaufmanniana; T. ferg – T. ferganica; T. tuberg – T. tubergeniana; T. fost – T. fosteriana; T. car – T. carinata; T. mog – T. mogoltavica.*

Random Forest and GBM gained the highest average results, performing excellent AUC (>0.999 both), TSS (>0.996 and >0.988 respectively) and Kappa (>0.982 and >0.97). MARS also performed excellent score by three metrics, but significantly lower than RF and GBM.

Our DNN model presented excellent average AUC (>0.969), TSS (>0.918) results in all pseudo-absence generating strategies, and good average Kappa (>0.829) score when using RSEB and RSEP pseudo-absence techniques.

### Pseudo-absence sampling strategy effect

Most SDMs showed increased prediction performance using RSEB and RSEP compared with RS. However, in some cases, RS performed better or as well as RSEB and RSEP techniques. RSEB gained the highest improvement of scores by all metrics for GLM in comparison with other strategies, whereas RSEP obtained the best results when performing by DNN, MARS, SRE, and MaxEnt models. CTA performed better when using RS, while GBM and RF achieved the highest results using all pseudo-absence selecting strategies without significant improvement under RSEB or RSEP. Table 5 compares the results of all models under RS, RSEB, and RSEP.

## Discussion

### Models comparison

Defining suitable area of distribution for rare species is crucial in conservational biology, management and ecology. SDMs provide simple way to determine habitat suitability for endangered species in comparison with traditional methods. However, there is not consistency in choosing the best SDM technique. Numerous researches were conducted to compare the efficiency of most popular SDMs: RF and MaxEnt in modeling rare species. Our results based on AUC, TSS, Cohen’s Kappa metrics illustrate superiority of RF and GBM methods when modeling rare Tulipa species in Uzbekistan. Our results agree with those of Mi (2017), which provides much higher performance of RF in cases with small sample sizes (<33) in comparison with MaxEnt.

Our simple DNN model performed with excellent average AUC and TSS results and good Kappa, but still was inferior to RF and GLM in most cases, however, it performed better than MaxEnt and other models. Few studies suggest improving DNN models by using Convolutional Neural Networks (Deneu, 2021), Bagging Ensembles (Rew, 2021), and Recurrent Neural Networks (Lee, 2018), and Transfer Learning (Rademaker et al. 2019). But DNN models are still required a large sample size of the dataset to perform better results than most common SDMs (Rademaker et al. 2019).

### Pseudo-absence generating strategy

Most widely used and popular SDMs require not only presence data but also absence data too. In most cases of defining suitable distribution area and finding new population of rare species in poor studied areas the real absence data is not available, so generating pseudo-absence data can be an alternative way to solve this problem. Random Sampling (RS) is the most widespread pseudo-absence method, it is simple and most SDMs use it by default. However, the efficiency of this method is not very high, so numerous researches suggested different alternatives (Iturbide et al. 2015).

We tested the effects of RS, RSEB, and RSEP pseudo-absence generating methods and found that RSEB and RSEP are able to gain better results than simple RS. Our results illustrate that RSEP is the best pseudo-absence strategy among studied techniques and this statement is in agreement with those of Rew (2021). It can significantly improve the results of most SDMs (DNN, GBM, RF, MARS, SRE, and MaxEnt), however, some models gained better performance under RS (CTA) or RSEB (GLM) pseudo-absence selecting strategy.

## Conclusion

By studying the performance of 9 different SDMs and 3 methods of pseudo-absence selecting strategies, we have found that using RF and GBM models when combining with RSEP technique is the best choice in studying species with small presence sample size and lack of real absence data (which is specific problem for rare endangered species in poor studied regions) in bioclimatic conditions of Uzbekistan.

## Supplementary materials

**S1**: Code and species distribution data; S2 Table: SDM performance using all methods for 8 rare *Tulipa* species

## Acknowledgements

No funding was received for conducting this study.

## Disclosure statement

No potential conflict of interest was reported by the authors.

## Notes

### Competing Interest Statement

The authors have declared no competing interest.

