## Supplementary materials for "Comparison of different models in predicting habitat suitability of rare species in Uzbekistan: 8 rare Tulipa species case-study"

**S1.** Code and species distribution data.

**1.** Our source code of pseudo-absence selecting using Python located in ‘preprocessing’ folder. It can be found in https://github.com/KhondamirRustamov/SDM-DL-ML-Tulipa-uz/tree/main/preprocessing.

**2.** Our source code of DNN modeling and training using Tensorflow Python package located in ‘SDMs/Python-DNN-model’ and can be found in https://github.com/KhondamirRustamov/SDM-DL-ML-Tulipa-uz/tree/main/SDMs/Python-DNN-model. Our source code GLM, GBM, RF, MARS, CTA, SRE, FDA, MaxEnt modeling using ‘BIOMOD2’ R package located in ‘SDMs/R’ and can be found in https://github.com/KhondamirRustamov/SDM-DL-ML-Tulipa-uz/tree/main/SDMs/R.

**3.** We upload all 8 species presence and pseudo-absence (RS, RSEB, RSEP) data in https://github.com/KhondamirRustamov/SDM-DL-ML-Tulipa-uz/tree/main/species_data.

**S2. Table:** SDM performance using all methods for 8 rare *Tulipa* species

| Species | Pseudo-absence selecting model | Metric | Modeling methods | | | | | | | | |
| --- | --- | --- | --- | --- | --- | --- | --- | --- | --- | --- | --- |
|  |  |  | **DNN** | **GLM** | **GBM** | **RF** | **MARS** | **CTA** | **SRE** | **FDA** | **MaxEnt** |
| *Tulipa kaufmanniana*  (n=45) | **RS** | **AUC** | 0.956 | 0.982 | 0.999 | **1.000** | 0.985 | 0.893 | 0.787 | 0.976 | 0.968 |
|  |  | **TSS** | 0.844 | 0.942 | 0.976 | **0.998** | 0.911 | 0.742 | 0.573 | 0.889 | 0.871 |
|  |  | **Kappa** | 0.734 | 0.773 | 0.976 | **0.988** | 0.800 | 0.741 | 0.610 | 0.760 | 0.841 |
|  | **RSEB** | **AUC** | 0.983 | 0.998 | **1.000** | **1.000** | **1.000** | 0.991 | 0.798 | 0.944 | 0.932 |
|  |  | **TSS** | 0.888 | 0.967 | 0.993 | **1.000** | **1.000** | 0.913 | 0.596 | 0.887 | 0.864 |
|  |  | **Kappa** | 0.935 | 0.937 | 0.988 | **1.000** | **1.000** | 0.878 | 0.709 | 0.924 | 0.910 |
|  | **RSEP** | **AUC** | 0.985 | 0.983 | **1.000** | **1.000** | 0.997 | 0.987 | 0.793 | 0.973 | 0.888 |
|  |  | **TSS** | 0.888 | 0.871 | 0.991 | **0.996** | 0.982 | 0.880 | 0.587 | 0.931 | 0.778 |
|  |  | **Kappa** | 0.935 | 0.840 | 0.964 | **0.976** | 0.928 | 0.889 | 0.667 | 0.915 | 0.864 |
| *Tulipa ferganica*  (n=37) | **RS** | **AUC** | 0.956 | 0.968 | **1.000** | **1.000** | 0.988 | 0.946 | 0.849 | 0.980 | 0.823 |
|  |  | **TSS** | 0.847 | 0.862 | **0.997** | **0.997** | 0.932 | 0.724 | 0.697 | 0.886 | 0.646 |
|  |  | **Kappa** | 0.803 | 0.686 | **0.985** | **0.985** | 0.782 | 0.802 | 0.681 | 0.728 | 0.756 |
|  | **RSEB** | **AUC** | 0.984 | 0.969 | **1.000** | **1.000** | 0.989 | 0.983 | 0.851 | 0.988 | 0.877 |
|  |  | **TSS** | 0.932 | 0.886 | 0.992 | **0.995** | 0.943 | 0.927 | 0.703 | 0.897 | 0.757 |
|  |  | **Kappa** | 0.729 | 0.668 | 0.970 | **0.985** | 0.830 | 0.874 | 0.703 | 0.792 | 0.850 |
|  | **RSEP** | **AUC** | 0.988 | 0.983 | 1.000 | **1.000** | 0.997 | 0.955 | 0.865 | 0.990 | 0.966 |
|  |  | **TSS** | 0.905 | 0.903 | 0.995 | **1.000** | 0.968 | 0.762 | 0.730 | 0.927 | 0.932 |
|  |  | **Kappa** | 0.651 | 0.826 | 0.985 | **1.000** | 0.925 | 0.816 | 0.831 | 0.848 | 0.705 |
| *Tulipa dubia*  (n=33) | **RS** | **AUC** | 0.963 | 0.987 | 0.999 | **1.000** | 0.921 | 0.955 | 0.842 | 0.993 | 0.877 |
|  |  | **TSS** | 0.893 | 0.939 | 0.988 | **0.997** | 0.842 | 0.836 | 0.685 | 0.979 | 0.755 |
|  |  | **Kappa** | 0.617 | 0.838 | 0.966 | **0.984** | 0.878 | 0.848 | 0.746 | 0.893 | 0.834 |
|  | **RSEB** | **AUC** | 0.980 | 0.993 | **1.000** | 0.999 | 0.998 | 0.929 | 0.845 | 0.982 | 0.969 |
|  |  | **TSS** | 0.954 | 0.976 | **0.997** | **0.997** | 0.985 | 0.812 | 0.691 | 0.948 | 0.939 |
|  |  | **Kappa** | 0.801 | 0.891 | **0.984** | **0.984** | 0.931 | 0.859 | 0.776 | 0.933 | 0.966 |
|  | **RSEP** | **AUC** | 0.976 | 0.997 | **1.000** | **1.000** | 0.997 | 0.949 | 0.845 | 0.968 | 0.938 |
|  |  | **TSS** | 0.905 | 0.958 | **1.000** | **1.000** | 0.955 | 0.830 | 0.691 | 0.921 | 0.876 |
|  |  | **Kappa** | 0.651 | 0.891 | **1.000** | **1.000** | 0.966 | 0.819 | 0.776 | 0.913 | 0.913 |
| *Tulipa affinis*  (n=19) | **RS** | **AUC** | 0.973 | 0.948 | 0.996 | **1.000** | 0.986 | 0.937 | 0.774 | 0.975 | 0.988 |
|  |  | **TSS** | 0.947 | 0.784 | 0.974 | **0.995** | 0.968 | 0.789 | 0.547 | 0.879 | 0.963 |
|  |  | **Kappa** | 0.774 | 0.595 | 0.907 | **0.972** | 0.859 | 0.744 | 0.575 | 0.779 | 0.826 |
|  | **RSEB** | **AUC** | **1.000** | 0.914 | 0.999 | **1.000** | 0.983 | 0.852 | 0.784 | 0.986 | 0.918 |
|  |  | **TSS** | **1.000** | 0.758 | 0.989 | **0.995** | 0.895 | 0.916 | 0.568 | 0.916 | 0.837 |
|  |  | **Kappa** | **1.000** | 0.514 | 0.945 | **0.972** | 0.826 | 0.981 | 0.663 | 0.826 | 0.878 |
|  | **RSEP** | **AUC** | **1.000** | 0.913 | **1.000** | **1.000** | **1.000** | 0.916 | 0.787 | 0.997 | 0.945 |
|  |  | **TSS** | **1.000** | 0.737 | **1.000** | **1.000** | **1.000** | 0.747 | 0.574 | 0.974 | 0.889 |
|  |  | **Kappa** | **1.000** | 0.471 | **1.000** | **1.000** | **1.000** | 0.757 | 0.688 | 0.889 | 0.911 |
| *Tulipa tubergeniana*  (n=15) | **RS** | **AUC** | 0.944 | 0.958 | 0.999 | **1.000** | 0.997 | 0.961 | 0.82 | 0.980 | 0.863 |
|  |  | **TSS** | 0.933 | 0.893 | 0.987 | **0.993** | 0.980 | 0.907 | 0.64 | 0.933 | 0.727 |
|  |  | **Kappa** | 0.717 | 0.625 | 0.962 | **0.964** | 0.899 | 0.832 | 0.66 | 0.804 | 0.799 |
|  | **RSEB** | **AUC** | 0.944 | **1.000** | **1.000** | **1.000** | 0.930 | 0.956 | 0.827 | 0.995 | 0.932 |
|  |  | **TSS** | 0.933 | **1.000** | **1.000** | 0.993 | 0.860 | 0.900 | 0.653 | 0.980 | 0.867 |
|  |  | **Kappa** | 0.717 | **1.000** | **1.000** | 0.964 | 0.887 | 0.804 | 0.718 | 0.899 | 0.922 |
|  | **RSEP** | **AUC** | 0.977 | 0.997 | 0.999 | **1.000** | **1.000** | 0.968 | 0.833 | 0.966 | 0.997 |
|  |  | **TSS** | 0.966 | 0.960 | 0.987 | **1.000** | **1.000** | 0.907 | 0.667 | 0.933 | 0.993 |
|  |  | **Kappa** | 0.840 | 0.962 | 0.962 | **1.000** | **1.000** | 0.922 | 0.784 | 0.962 | 0.964 |
| *Tulipa fosteriana*  (n=12) | **RS** | **AUC** | **1.000** | **1.000** | **1.000** | **1.000** | **1.000** | 0.983 | 0.708 | **1.000** | 0.958 |
|  |  | **TSS** | **1.000** | **1.000** | **1.000** | **1.000** | **1.000** | 0.967 | 0.417 | **1.000** | 0.917 |
|  |  | **Kappa** | **1.000** | **1.000** | **1.000** | **1.000** | **1.000** | 0.841 | 0.565 | **1.000** | 0.952 |
|  | **RSEB** | **AUC** | 0.930 | **1.000** | **1.000** | **1.000** | **1.000** | 0.986 | 0.708 | **1.000** | 0.996 |
|  |  | **TSS** | 0.875 | **1.000** | **1.000** | **1.000** | **1.000** | 0.900 | 0.417 | **1.000** | 0.992 |
|  |  | **Kappa** | 0.608 | **1.000** | **1.000** | **1.000** | **1.000** | 0.867 | 0.565 | **1.000** | 0.956 |
|  | **RSEP** | **AUC** | **1.000** | **1.000** | **1.000** | 0.999 | **1.000** | 0.922 | 0.704 | 0.995 | 0.871 |
|  |  | **TSS** | **1.000** | **1.000** | **1.000** | 0.992 | **1.000** | 0.825 | 0.408 | 0.992 | 0.742 |
|  |  | **Kappa** | **1.000** | **1.000** | **1.000** | 0.956 | **1.000** | 0.857 | 0.527 | 0.956 | 0.802 |
| *Tulipa carinata*  (n=17) | **RS** | **AUC** | 0.985 | 0.988 | 0.999 | **1.000** | **1.000** | 0.938 | 0.838 | 0.964 | 0.968 |
|  |  | **TSS** | 0.941 | 0.906 | 0.982 | 0.994 | **1.000** | 0.859 | 0.676 | 0.929 | 0.935 |
|  |  | **Kappa** | 0.771 | 0.900 | 0.967 | 0.968 | **1.000** | 0.816 | 0.676 | 0.905 | 0.935 |
|  | **RSEB** | **AUC** | **1.000** | **1.000** | **1.000** | **1.000** | **1.000** | 0.929 | 0.844 | 0.969 | 0.879 |
|  |  | **TSS** | **1.000** | **1.000** | 0.994 | **1.000** | **1.000** | 0.812 | 0.688 | 0.929 | 0.759 |
|  |  | **Kappa** | **1.000** | **1.000** | 0.968 | **1.000** | **1.000** | 0.834 | 0.727 | 0.905 | 0.824 |
|  | **RSEP** | **AUC** | **1.000** | 0.980 | **1.000** | **1.000** | **1.000** | 0.959 | 0.844 | 0.970 | 0.970 |
|  |  | **TSS** | **1.000** | 0.876 | 0.994 | **1.000** | **1.000** | 0.841 | 0.688 | 0.929 | 0.941 |
|  |  | **Kappa** | **1.000** | 0.767 | 0.968 | **1.000** | **1.000** | 0.863 | 0.727 | 0.932 | 0.967 |
| *Tulipa mogoltavica*  (n=15) | **RS** | **AUC** | 0.977 | **1.000** | **1.000** | **1.000** | **1.000** | 0.948 | 0.697 | 0.998 | 0.930 |
|  |  | **TSS** | 0.933 | **1.000** | **1.000** | **1.000** | **1.000** | 0.780 | 0.393 | 0.987 | 0.860 |
|  |  | **Kappa** | 0.717 | **1.000** | **1.000** | **1.000** | **1.000** | 0.780 | 0.518 | 0.931 | 0.887 |
|  | **RSEB** | **AUC** | 0.988 | **1.000** | **1.000** | **1.000** | **1.000** | 0.690 | 0.697 | **1.000** | 0.997 |
|  |  | **TSS** | 0.966 | **1.000** | **1.000** | **1.000** | **1.000** | 0.380 | 0.393 | **1.000** | 0.993 |
|  |  | **Kappa** | 0.840 | **1.000** | **1.000** | **1.000** | **1.000** | 0.463 | 0.518 | **1.000** | 0.964 |
|  | **RSEP** | **AUC** | **1.000** | **1.000** | **1.000** | **1.000** | **1.000** | 0.690 | 0.700 | **1.000** | 0.997 |
|  |  | **TSS** | **1.000** | **1.000** | **1.000** | **1.000** | **1.000** | 0.380 | 0.400 | **1.000** | 0.993 |
|  |  | **Kappa** | **1.000** | **1.000** | **1.000** | **1.000** | **1.000** | 0.463 | 0.548 | **1.000** | 0.964 |
